# Transient interactions between the fuzzy coat and the cross-β core of brain-derived Aβ42 filaments

**DOI:** 10.1101/2024.01.08.574772

**Authors:** Maria Milanesi, Z. Faidon Brotzakis, Michele Vendruscolo

## Abstract

A wide range of human disorders, including Alzheimer’s disease (AD), are characterised by the aberrant formation of amyloid fibrils. Amyloid fibrils are filamentous structures characterized by the presence of a highly-ordered cross-β core. In many cases, this core structure is flanked by disordered regions, often referred to as fuzzy coat. The structural properties of fuzzy coats, and the way in which they interact with their environments, however, have not been described in full detail to date. Here, we generated the conformational ensembles of two brain-derived amyloid filaments of Aβ42, corresponding respectively to familial and sporadic forms of AD. The approach that we used, called metadynamic electron microscopy metainference (MEMMI), enabled us to provide a characterization of the transient interactions between the fuzzy coat and the cross-β core of the filaments. These calculations indicated that the familial AD filaments are less soluble than the sporadic AD filaments, and that the fuzzy coat contributes to increasing the solubility of both types of filament. In addition, by analyzing the deviations between the density maps from cryo-EM and from the MEMMI structural ensembles, we observed a slowing down in the diffusion of water and sodium ions near the surface of the filaments, offering insight into the hydration dynamics of amyloid fibrils. These results illustrate how the metainference approach can help analyse cryo-EM maps for the characterisation of the properties of amyloid fibrils.

## Introduction

Alzheimer’s disease (AD) causes cognitive impairment through neuronal loss, and accounts for 60-80% of the cases of dementia worldwide (1, 2). The accumulation of intracellular neurofibrillary tangles, formed by the tau protein, and of extracellular senile plaques, formed by the amyloid β peptide (Aβ), represents the main hallmark of the disease at the molecular level (3-5). In the amyloidogenic pathway, Aβ is generated through successive cleavages of the amyloid precursor protein (APP) by β-secretase and β-secretase (5). Aβ is poorly soluble and readily aggregates into amyloid fibrils (6). The surface of the amyloid fibrils acts as a catalyst for secondary nucleation, with a feedback mechanism that facilitates the proliferation of the amyloid fibrils themselves (6, 7).

The structures of amyloid fibrils are characterised by a distinctive molecular architecture known as cross-β (6, 8). This architecture consists of β strands that stack perpendicularly to the long axis of the fibril, forming structures called protofilaments. Multiple protofilaments can then associate laterally, giving rise to twisted fibrils (6, 8, 9). Investigations of synthetic Aβ peptides in in vitro preparations revealed a spectrum of different polymorphic structures of Aβ fibrils (10-14). Furthermore, recent reports indicated that these fibrils are structurally different from amyloid fibrils derived from brain tissue (15-17). The cryo-EM structures of Aβ42 fibrils extracted from the parenchyma brain tissue of 10 AD patients were recently reported (17). Two distinct polymorphs were distinguished, referred to as, respectively, type I filaments, principally found in sporadic AD (sAD), and type II filaments, present in familial AD (fAD) and other conditions. The atomic model of the type I filament spans from residues G9 to A42, forming the organized structural core of the protofilament structure, which exhibits the characteristic cross-β architecture of amyloid fibrils. This type I filament consists of two S-shaped protofilaments closely packed together through hydrophobic interactions, exhibiting pseudo-2_1_ symmetry. In contrast, the cross-β core of the type II filament spans from residues V12 to A42, and it is made up of two protofilaments forming the fibril with a C2 symmetry. The structural models obtained from both filaments are missing about 20% of the Aβ42 sequence, represented by the N-terminal residues. This N-terminal regions is expected to remain disordered, displaying a high conformational heterogeneity, collectively forming a fuzzy coat around the cross-β core of the filaments (18-21). Further analysis of the structural properties of the disordered regions in the amyloid fibrils requires methods capable of characterizing conformational heterogeneity. The used of cryo-EM for this purpose remains quite challenging (22-31). Consequently, most structural studies on amyloid fibrils primarily concentrate on characterizing conformations and interactions within the cross-β cores, while the more dynamic flanking regions remain less explored.

The prominent role of the N-terminal region of Aβ42 fibrils in the aggregation process and the stability of the amyloid fibrils has been documented in several studies. This is substantiated by the observed influence of various N-terminal mutants on the aggregation process and the subsequent neurotoxic effects of the peptide (32-36). Nevertheless, our understanding of the impact of the fuzzy coat region on various functionalities and characteristics of the amyloid filaments, such as its interactions with the cross-β core, its contribution to the binding between the amyloid filaments and other biologically relevant components, and the solubility of the amyloid filaments, remains elusive. These insights would be of great importance for pharmacological research, as they could guide the development of new drugs and deepen our understanding of the molecular mechanisms of existing drugs targeting the N-terminal region of the filaments.

In this study, we report the structural ensembles of Aβ42 in the type I and type II filaments, obtained using the metadynamic electron microscopy metainference (MEMMI) method (37). The analysis of these structural ensembles provides insights into the role of the fuzzy coat in modulating the properties of amyloid fibrils.

## Results

### Structure and dynamics of type I and type II filaments

The structures of the cross-β cores (residues 9-42) of Aβ42 filaments derived from AD brains were recently reported by cryo-EM (17). These cross-β cores exhibit two types of structurally related protofilaments, resulting in the formation of two distinct filament types. Type I filaments were predominantly found in the brains of individuals with sAD (PDB: 7q4b, EMD-15770), whereas type II filaments (PDB: 7q4m, EMD-15771) were primarily observed in the brains of individuals with fAD and other misfolding-related pathologies. In this study, we determine the structural ensemble of the whole Aβ42 filaments (residues 1-42) using the MEMMI method (37). Multiple collective variables (CVs) were biased using metadynamics (38, 39) to accelerate the conformational sampling. Diffusion traces of the CVs and the free energy profiles as a function of simulation time of all CVs indicate that the simulations are converged (**Figures S1** and **S2)**. We modelled the system as a stack of twelve peptides on each side. Within the cross-β core, a parallel β-sheet structure is consistently maintained. By contrast, we observed significant conformational heterogeneity in the N-terminal (residues 1-9) for both type I (**Figure 1A,C**) and type II (**Figure 1B,D**) filaments. The resulting conformational ensembles can be described as a fuzzy coat that surrounds the cross-β core of the filaments.

**Figure 1.**
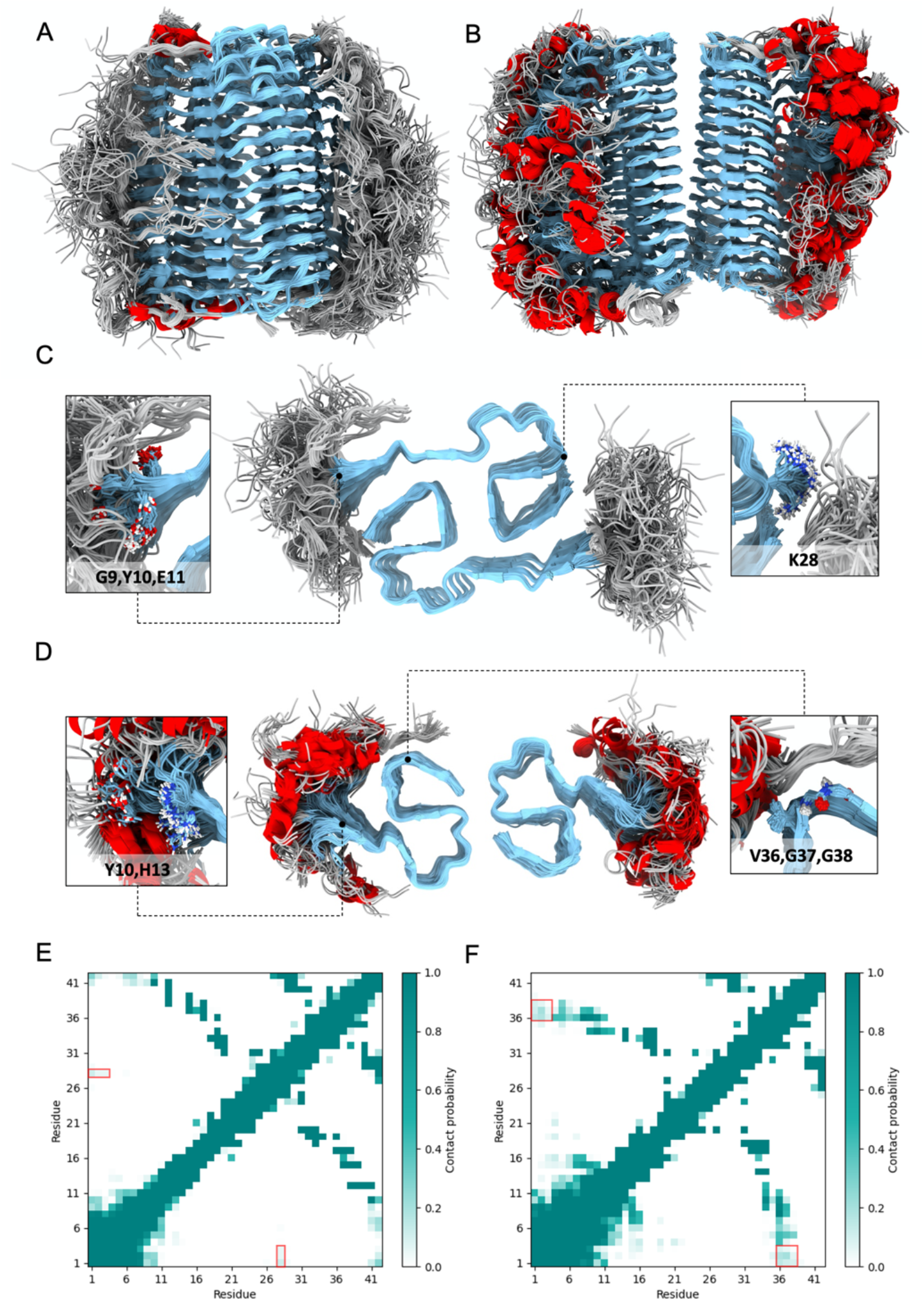
Structural ensembles of type I and type II filaments formed by Aβ42. **(A,B)** Structural ensembles of type I (A) and type II (B) filaments, obtained by extracting 100 conformations from the structural ensembles generated in this work. **(C**,**D)** Lateral view of the structural ensembles and close-up of the residues shielded by the fuzzy coat of type I (C) and type II (D) filaments. The colors denote secondary structure elements: β-sheet (cyan), coil (grey), and *α*-helix (red). The insets show residues G9, Y10, E11, and K28 in type I filaments, and residues Y10, H13, V36, G37, and G38 in type II filaments. These residues are on the surface of the cross-β core, but are partially shielded from the solvent by interactions with the fuzzy coat. **(E**,**F)** Contact maps corresponding to the structural ensembles of type I (E) and type II (F) filaments. The contact maps show the contact probability between residue pairs. Contacts with probability 1 (teal) are always formed, contacts with probability 0 (white) are never formed, and contacts with intermediate probabilities are transient. As examples the red boxes highlight the transient contacts between residues D1, A2 and E3 with residue K28 (E) and between residues D1, A2 and E3 with residues V36, G37 and G38 (F).

### Fuzzy coat interactions with the cross-β core

The fuzzy coat created by these N-terminal disordered regions extends to make contacts with the cross-β core. To investigate these contacts, we determined that the average distance between the fuzzy coat and the cross-β core, as quantified by the distribution of the CV dcm (**Table S2**), is approximately 3 nm for both filament types (**Figure S3**). To gain insights into how the fuzzy coat affects the structure and dynamics of the filaments, we performed a solvent-accessible surface area (SASA) analysis (**Figure 3**). Comparing the SASA per residue of the structural ensemble of the complete filament structure with the SASA obtained when excluding the N-terminus residues 1-9, we observed a SASA reduction for residues G9, Y10, E11, and K28 in type I filament, and residues Y10, H13, V36, G37, and G38 for type II filament. This analysis identified the specific residues that are shielded by the presence of the fuzzy coat during the filament dynamics (**Figures 1C,D** and **3C,D**).

### Secondary structure of the fuzzy coat and cross-β core

Within the cross-β core, our findings support the β-sheet composition originally reported (17). Our calculations reveal an additional transient short β-motif at residues 24-25 in the type I filament **(figure 2A**, left panel). Moreover, residues 30-36, combining the β3 and β4 regions (17), form a single β-motif (**Figure 2A**). Similar trends are also observed in the type II filament (**Figure 2B**). Furthermore, our results indicate a higher *α*-helical content in the fuzzy coat of the type II filament (residues 2-8) compared to the type I filament, which exhibits a more coiled and disordered structure. These findings align with similar observations made in other amyloidogenic proteins, such as huntingtin, where the fuzzy coat adopts an *α*-helical structure, suggesting potential biologically relevant functions (40).

**Figure 2.**
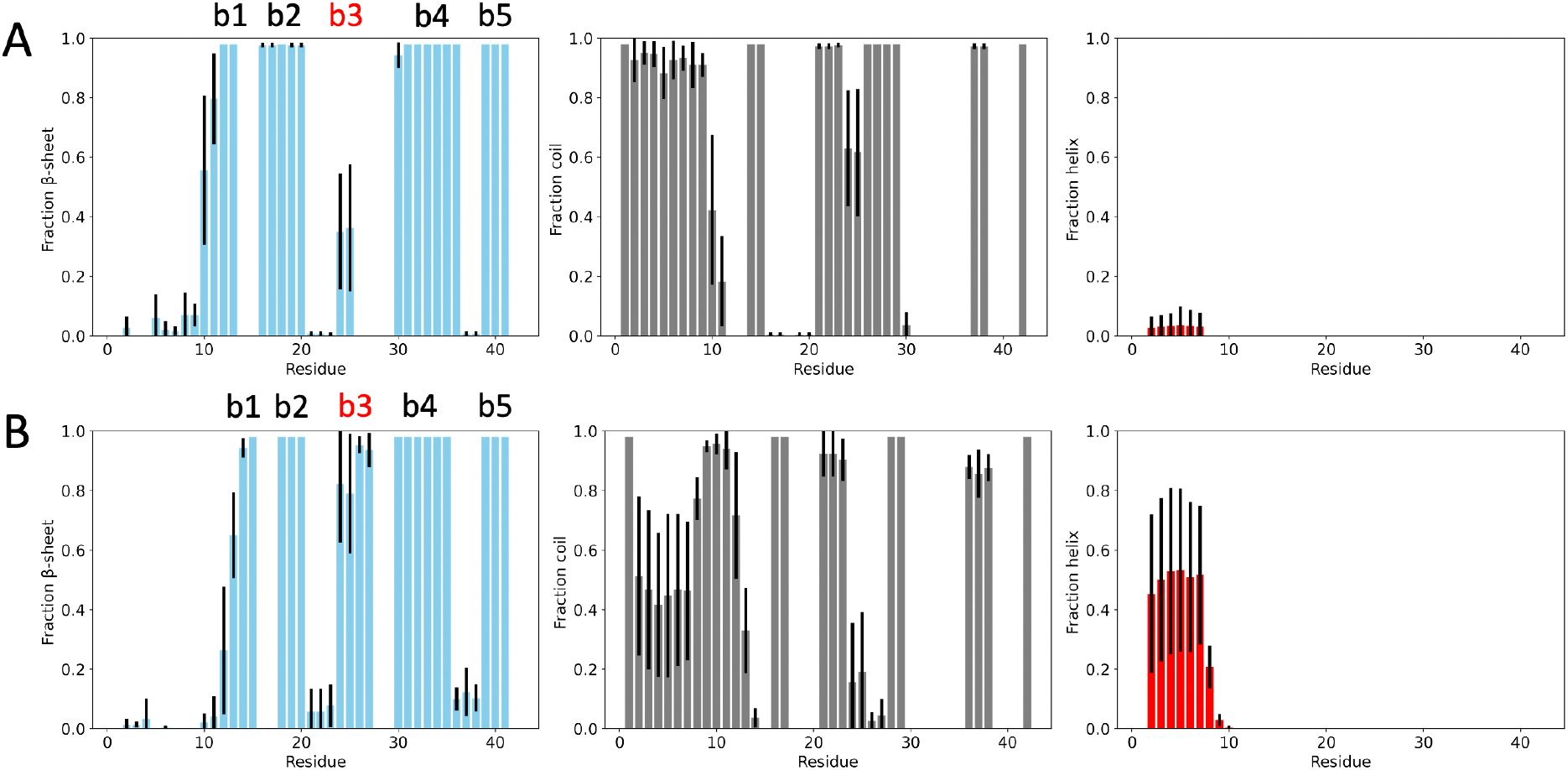
Secondary structure populations of the structural ensembles of type I and type II filaments. **(A**,**B)** Type I filaments (A) and type II filaments (B). Error bars are calculated as standard deviations between the first and second halves of the MEMMI simulations. The colors are based on the secondary structure elements: β-sheet (cyan), coil (grey), and *α*-helix (red).

**Figure 3.**
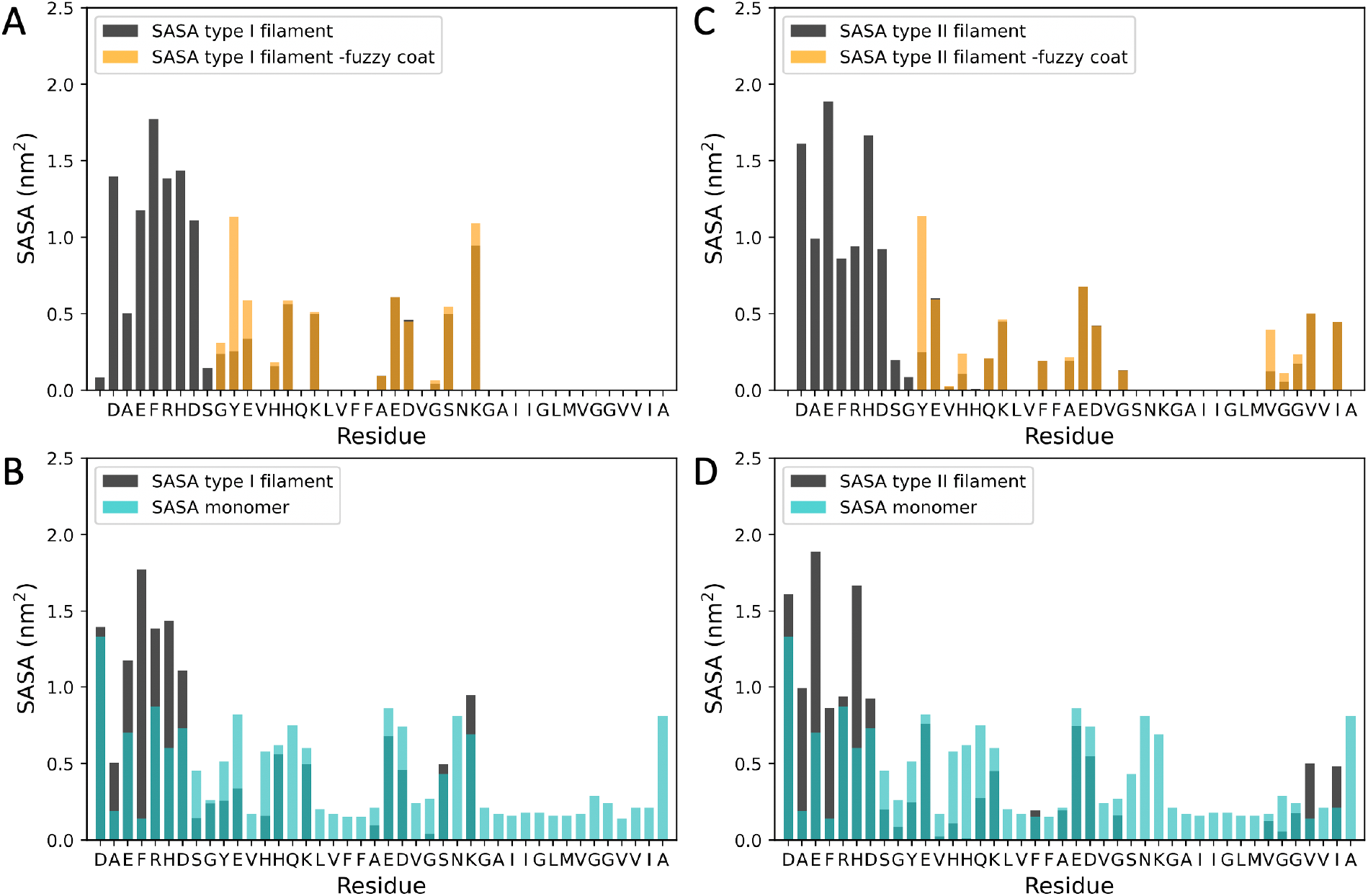
Effect of the fuzzy coat on the solvent-exposed surface area (SASA) for type I and type filaments. **(A-D)** Black bars represent the per residue SASA analysis of type I filament (A,B) and type II filament (C,D). These SASA values were calculated based on 100 conformations extracted from the MEMMI structural ensemble. Orange bars in (A, C) show the per residue SASA values of the 100 obtained structures after removing the N-terminus. Light blue bars in (B, D) correspond to SASA values of the Aβ42 monomers in solution (41).

### Influence of the fuzzy coat on the solubility scores of type I and type II filaments

The comparison of the SASA of the whole structure of the two Aβ42 filaments types with respect to the SASA of the Aβ42 monomer in solution (41) indicates that the hydrophobic region in the C-terminus region of the protein (37-42) is not exposed to the solvent in type I filaments (**Figure 3A**), generating a more soluble filament (**Figure 4A**), according to the structurally-corrected CamSol solubility score (42, 43). Instead in the type II filaments it remains exposed to the solvent (**Figure 3B**), generating a more insoluble filament (**Figure 4B**, red box). This result is also confirmed by the electrostatic potential surface of the two filament types (**Figure S4**).

**Figure 4.**
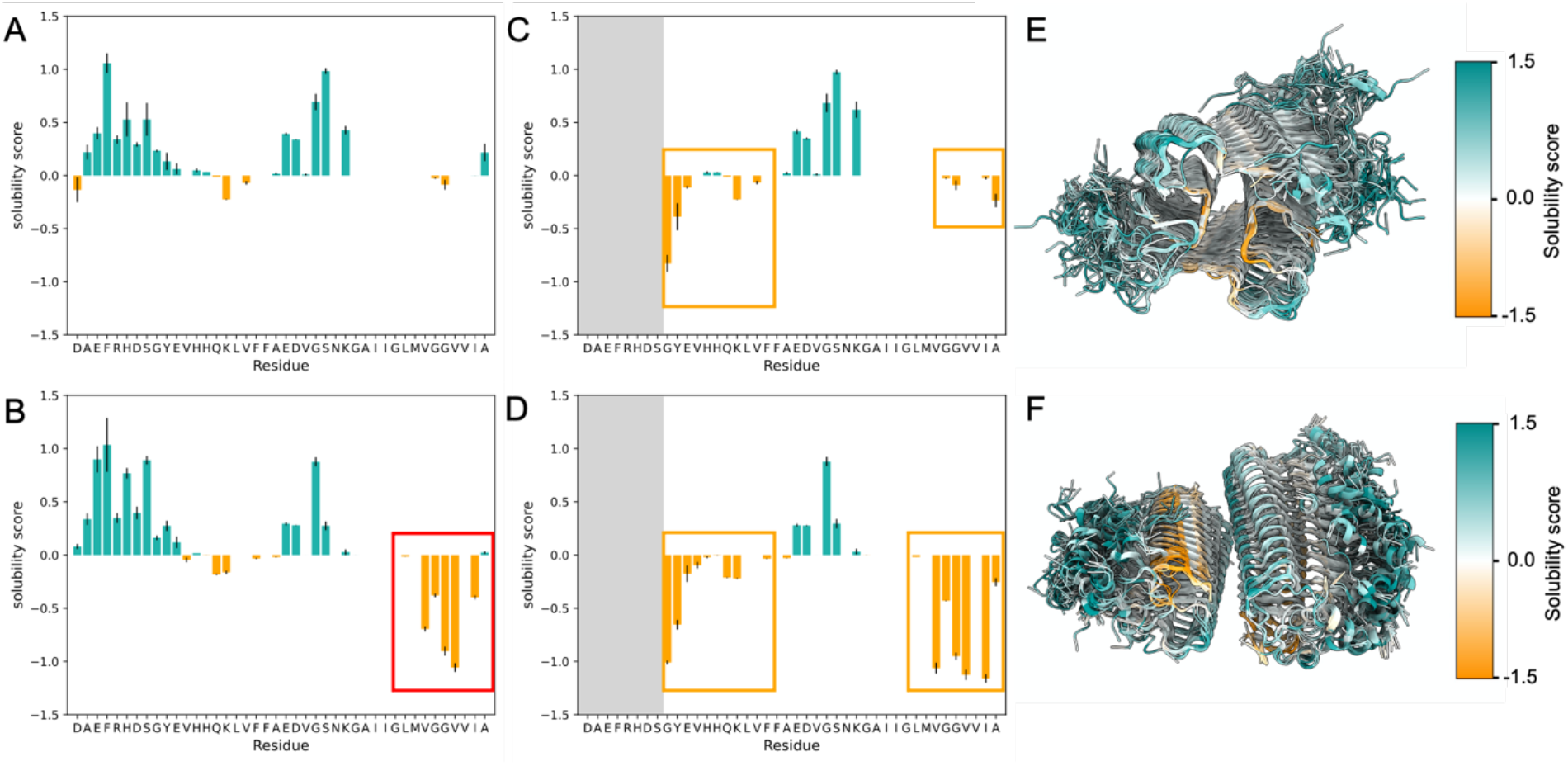
Influence of the fuzzy coat on the solubility scores of type I and type II filaments. **(A**,**B)** CamSol structurally-corrected solubility score per residue for type I (A) and type II (B) filaments; mean and standard deviations are calculated from 20 randomly selected conformations from the MEMMI structural ensemble. Residues with solubility scores above 0 tend to be soluble, shown in light sea-green, while negative values indicate insoluble residues, shown in orange. The red square indicates the region of low solubility score in type II filaments. In type I filaments, this region is protected because it is buried in the core of the amyloid structure. **(C**,**D)** CamSol structurally-corrected solubility scores per residue for type I (C) and type II (D) filaments; the same 20 conformations analysed in (A,B) are shown, except that the fuzzy coat was removed. The orange squares indicate regions of low solubility score that are partially protected by the fuzzy coat. These conformations correspond to type I (A,C) and type II (B,D) filaments. **(E**,**F)** Graphical representation of the 20 conformations colored by the solubility score, with orange indicating less soluble and light sea-green indicating more soluble regions.

To explore the effects of the fuzzy coat on the solubility of the cross-β core, we calculated the structurally-corrected CamSol solubility score obtained by excluding the fuzzy coat (**Figure 4C-F**). We calculated the average CamSol solubility score for the residues in the cross-β core for the type I and type II filaments. This calculation was performed for both the ensemble with the fuzzy coat present and the ensemble from which the fuzzy coat was removed (**Table 1**). The comparison of these solubility scores indicates that the fuzzy coat increases the solubility scores of the cross-β core of the filaments (**Figure 3C,D**, orange squares). This result is in accordance with the shielded residues observed in the SASA analysis reported above.

**Table 1.** Average solubility values with corresponding errors for the cross-β core of type I and type II Filaments. This table presents the averages of solubility values, along with their corresponding errors, for the cross-β core of type I and type II filaments. The values are derived from a 20-conformation ensemble, considering both ensembles with (+ fuzzy coat) and without (- fuzzy coat) the fuzzy coat.

|  | Type I Filament | Type II Filament |
| --- | --- | --- |
| + fuzzy coat | 0.02 ± 0.01 | -0.48 ± 0.07 |
| - fuzzy coat | -0.05± 0.08 | -0.71 ± 0.09 |

These calculations indicate that the filaments observed in familial AD (i.e. type II filaments) are less soluble than the filaments observed in sporadic AD (i.e. type II filaments), and that for both filaments the fuzzy coat may play a relevant role in increasing the solubility of the filaments.

### Correlation between experimental and calculated cryo-EM maps

To assess the fidelity of the MEMMI structural ensembles in capturing the molecular behaviour and dynamics of type I and type II filaments, we examined their correlation with experimental cryo-EM maps (**Figure 5**). The structural ensembles derived from type I (**Figure 5A,B**) and type II (**Figure 5E,F**) filaments exhibited generally a better correlation with their respective experimental cryo-EM maps compared to the previously reported single structures (PDB: 7Q4B and 7Q4M). The coefficients of correlation for the structural ensembles with the experimental density map were 0.83 for type I filament and 0.82 for type II filament. For comparison, the correlation coefficients for the single structures were 0.65 and 0.70, respectively. These results underscore the enhanced accuracy achieved by fitting structural ensembles to the cryo-EM maps. Furthermore, an essential feature of MEMMI is its capability to estimate the error in the experimental density map **(Figures 5C,D,G,H)**. The average relative error per Gaussian data point was determined to be 0.2 for both filaments. Here, the relative error represents the deviation of each Gaussian data point concerning the total overlap between all data Gaussian Mixture Model (GMM) and the specific component of the data GMM (see Equation 1).

**Figure 5.**
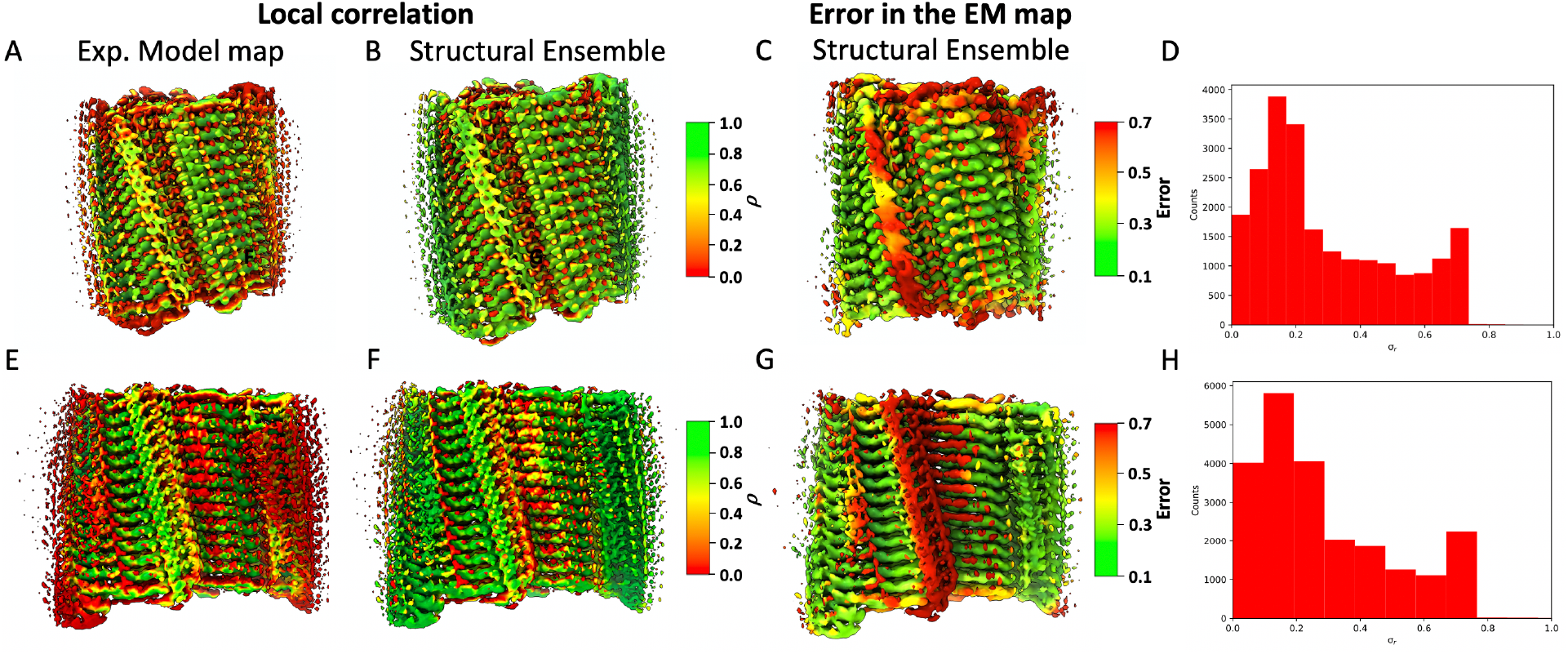
Local correlations and errors in the data from the MEMMI simulations. **(A-H)** Assessment of the local correlation between the cryo-EM map EMD-13800 (A,B) and EMD-13809 (E,F) with the cryo-EM maps generated by a previously determined single structure PDB:7Q4B (A), PDB:7Q4M (E), and the respective MEMMI structural ensemble (B,F). Map projection (C,G) and histogram (D,H) of the error in the GMM data for the type I filament (upper panel) and type II filament (lower panel) as obtained by the respective structural ensemble

### Ordered behaviour of solvent molecules

The calculated correlation and error maps indicate discrepancies in specific regions of the experimental density map. These regions are not effectively captured by either the single structure or the MEMMI structural ensembles, resulting in relatively low correlation and high error values (**Figure 5C,G)**. In both type I and type II filaments, these regions concern low-density volume data present over a groove formed around residues H14 and L21 by the filament structure (type I filaments), and at the interface around residues I41 and S26 of the two protofilaments (type II filaments). Additionally, another volume with the same characteristics is observed over residues E22 and D23 in both filaments.

A close-up examination of these regions reveals that the discrepancy is not associated to the density of the residues of the peptide **(Figure 6A)**. As previously highlighted (17), there are extra densities in the experimental density maps that were not utilized in modelling the atomic structure of the peptide. These additional densities may originate from solvent molecules with slower dynamics. To verify the hypothesis that solvation layer solvent molecules are trapped in these regions, exhibiting slower dynamics and therefore contributing to the experimental density map, we performed a solvent diffusion analysis for water and ions molecules during the MEMMI simulation **(Figure 6B-D)**. Notably, for both the filaments types, the oxygen atoms of water molecules moving within the grooves exhibit a lower diffusion coefficient compared to the diffusion coefficient of the bulk water molecules in the simulation box, as well as to water molecules near another point on the protein surface used as a control (around residues E11 for type I filament and G37 respectively, **Figure 6C**). These findings support the conclusion that water molecules diffuse slower within these grooves, as a result of interactions with the protein, as found in different studies (44).

**Figure 6.**
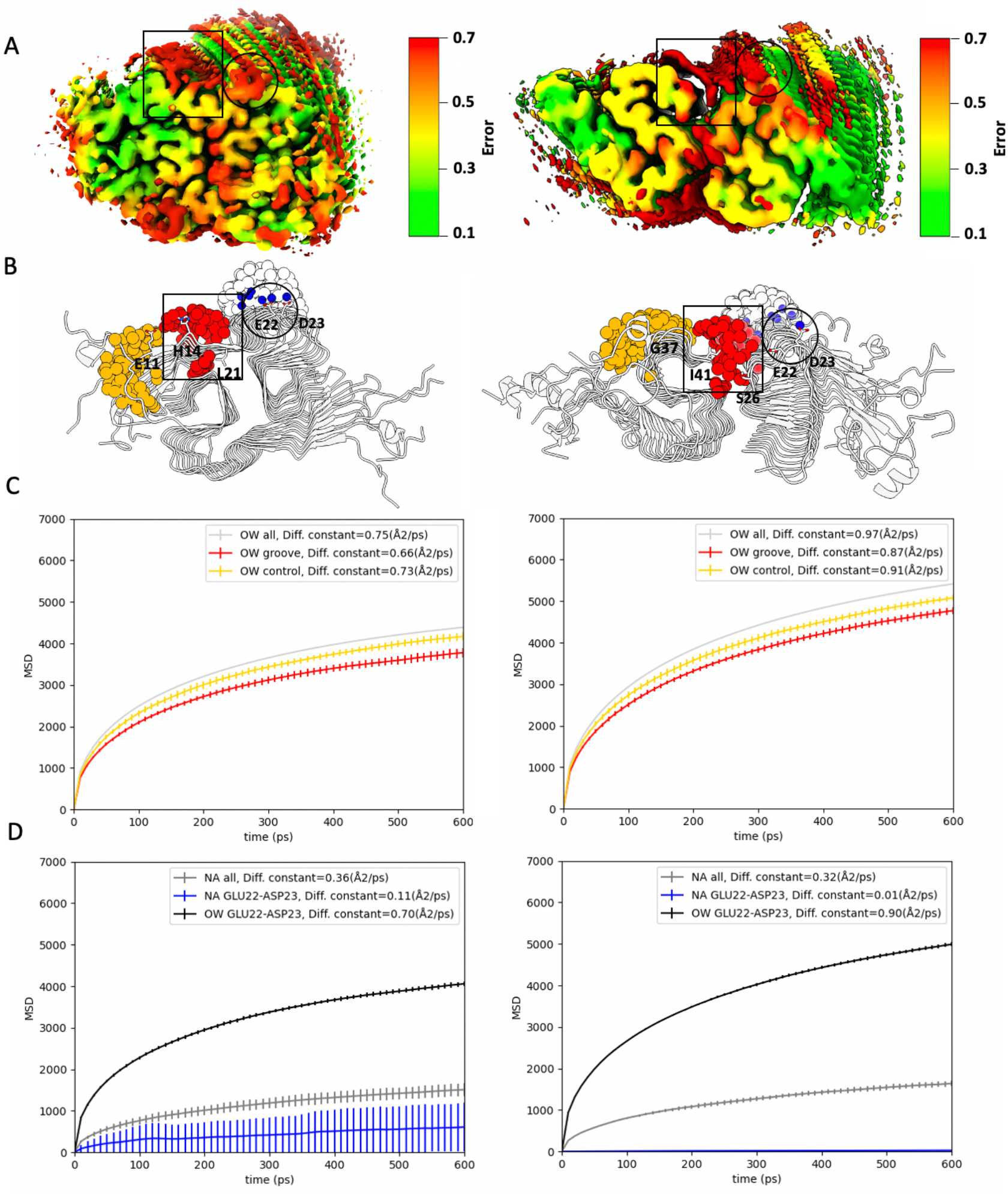
Analysis of the diffusion of water and ions near the surface of type I and type II filaments. **(A)** Lateral view of the error map of type I (left) and type II (right) filaments; black squares indicate the high error volumes of interest. **(B)** Schematic representation of selected atoms for the calculation of the diffusion coefficient of water molecules and sodium ions. Red spheres denote oxygen atoms of water molecules moving within the groove defined by residues H14 and L21 for type I filament, and by residues I41 and S26 for type II filament. Yellow spheres represent oxygen atoms of water molecules near residue E11 for type I filament and G37 for type II filament, used as controls. Blue spheres represent sodium atoms selected over residues E22 and D23. Transparent spheres highlight oxygen atoms of water molecules used as controls in comparison to sodium atoms. **(C,D)** The mean square displacement (MSD) analysis conducted on three replicas with the associated standard deviations are plotted over time for the water molecules (C) and sodium ions (D) experiments. Lines are color coded as in the schematic representation (B), with the exception of oxygen atom, shown in light grey, and all sodium ions, shown in grey, present in the simulation box and not shown in (B). In the legend, the diffusion constant for each selection is indicated.

Furthermore, the densities observed in correspondence of residues E22 and D23 have been linked to the capability of these two residues to coordinate positive ions (17). Consequently, we examined the diffusion of sodium ions. We found that the diffusion coefficient of the sodium ions moving over these two residues is lower than the diffusion coefficient of bulk sodium ions present in the simulated system (**figure 6D**). This observation suggests that sodium ions bind transiently residues E22 and D23 in both filament types. Overall, these findings demonstrate how the water dynamics is influenced by the presence of the filaments. The characterization of this interplay provides an illustration of the complex influence that amyloid fibrils have on their environments.

### Type I and type II filaments interact with aducanumab

The MEMMI structural ensembles provide valuable insights about the interactions of the amyloid fibrils with other biologically relevant components. Recently, the antibody aducanumab was approved by the Food and Drug Administration (FDA) for clinical use to treat Alzheimer’s disease (45, 46). Aducanumab binds the monomeric form as well as the oligomeric and fibrillar forms of Aβ42. It was shown in particular that aducanumab has a high affinity for the N-terminal region of Aβ42 (47). However, although aducanumab is able to decrease secondary nucleation (48), the mechanism by which this is achieved has not been clarified. Towards this goal, we report a schematic representation of a possible complex formed by aducanumab and Aβ42 filaments, which illustrates how one arm of the antibody binds the fuzzy coat, the other arm by shield the cross-β core, thus blocking potential secondary nucleation sites (**Figure S5**).

### Conclusions

We have reported the conformational ensembles of two brain-derived amyloid filaments of Aβ42, which provide a description of their highly dynamical fuzzy coats. The analysis of these structural ensembles has enabled us to illustrate how the fuzzy coats interact with the cross-β cores of the filaments, thus influencing their accessibility to the environment and in turn the solubility of the filaments themselves. Since the Bayesian approach adopted in the MEMMI method used here enables the determination of conformational ensembles consistent with the cryo-EM density maps, while also estimating the errors resulting from the procedure, we have been able to characterise the slowing down of the diffusion of water molecules and sodium ions near the surface of the amyloid filaments. Based on these results, we suggest that metainference approach described here will help analyse cryo-EM maps for the characterisation of the properties of heterogeneous macromolecular systems.

## Methods

### The MEMMI method

MEMMI (37) is a Bayesian method that models statistical ensembles of biomolecules by combining cryo-EM experimental data and metadynamics molecular dynamics in order to infer an ensemble of structures that maximally agree with the cryo-EM data, while simultaneously inferring various sources of errors (e.g. data error, forward model error, limited number of replicas able to model the heterogeneity of the system) (38). In MEMMI, the configurations of the structural ensemble and the different types of error are sampled from the following hybrid energy function

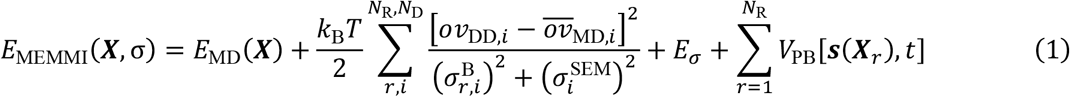

where *ϕ*_*D*_(***x***) is Data Gaussian Mixture model (GMM) conversion of the cryo-EM voxel map data to consisting of *N*_D_ Gaussian components, *ϕ*_*M*_ (***x***) is the model GMM, a GMM conversion of the molecular dynamics models, *ov*_*MD,i*_ is the overlap function quantifying the agreement between models generated by molecular dynamics (MD) and the data GMM is calculated by the following overlap function *ov*_MD, *i*_, *N*_R_ is the number of replicas of the system, dealing with the heterogeneity of the system, 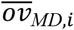 is the overlap between model GMM and data GMM is estimated over the ensemble of replicas as an average overlap per GMM component *ov*_*MD, i*_, 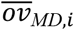 is the data GMM self-overlap per datapoint, **X** is a vector representing the atomic coordinates of the full ensemble, consisting of individual replicas 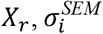 is the error incurred by the limited number of replicas in the ensemble, per datapoint., 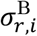 is the error due to random and systematic errors in the prior, forward model, and experiment per datapoint and replica, *E*_*σ*_ is the error energy associated with the error *σ* = (*σ*^*B*^, *σ*^*SEM*^) (see Ref. 40), *E*_MD_ is the force field, *s*(***X***) is the CV set (**Tables S1** and **S2**), and *V*_PB_ is the time-dependent biasing potential acting on a set of *N*_CV_ CVs (49). MEMMI samples the space of conformations *X*_*r*_ by multi-replica molecular dynamics simulations, while the error parameters for each datapoint 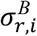 are sampled by a Monte Carlo sampling scheme at each time step.

### Initial filament structure

We build the initial structure of the full length filament by starting from the PDB databank deposited type-I (PDB: 7Q4B) and type-II (PDB: 7Q4M) Aβ filament structures, which only contains the cross-β core of the filament and a five chains stacked on top of each other forming a β-sheet, followed by extending the filament to a twelve chain stack by using Chimera (50) and the symmetry of each cross-β core (helical C1 and C2 symmetry respectively) in order to avoid finite size effects in the generated structural ensemble. Then, by using RosettaFold (51, 52) we model each the full-length polymorph filament by using each 12-stack cross-β core as a template and by providing the Aβ sequence. The cryo-EM density maps used as data in MEMMI are EMD-13800 and EMD-13809.

### Molecular dynamics setup and equilibration

For each full-length polymorph structure (type-I and type-II), we continue by creating a 11.99 × 7.40 × 9.51 and 9.30 × 13.68 × 9.56 nm simulation box, solvating with 22,334 and 35,090 water molecules, respectively, and neutralizing the net charge by adding 72 Na^+^ ions. We use the AMBER99SB-ILDN (53) force field and TIP3P (54) water models. We continued with an energy minimization followed by a 500 ps NPT equilibration at a temperature of 310 K and pressure of 1 atm, followed by an additional 2 ns NVT equilibration at 310 K. The molecular dynamics parameters are the same used previously (30, 31).

### MEMMI simulations

We first express the experimental voxel map data as a data GMM containing 22,492 and 22,496 Gaussians in total, resulting in a 0.878 and 0.885 correlation to the original voxel maps EMD-13800 and EMD-13809, using the gmmconvert utility (55). We continued by extracting 8 equally distant configurations throughout the 2 ns simulations from the previous NVT equilibration step and start MEMMI simulations, consisting of 8 replicas, resulting in an aggregate runtime of 1 μs, using PLUMED (56) (PLUMED.2.6.0-dev) and GROMACS (57) (GROMACS-2020.5). MEMMI is conducted in the NVT ensemble at 310 K using the same parameters as in the equilibration step. Configurations are saved every 10 ps for post-processing. The cryo-EM restraint is updated every 2 steps, using neighbour lists to compute the overlaps between model and data GMMs, with a neighbour list cut-off of 0.01 and update frequency stride of 100 steps. The CVs used for biasing in the simulations are shown in **Tables S1** and **S2**, and the biasing scheme is PBMetaD (49) with the well-tempered (58) and multiple-walkers (59) protocols. The hill height is set to 0.3 kJ/mol, with a deposition frequency of 200 steps and an adaptive Gaussians diffusion scheme (60). The biasing CVs correspond to degrees of freedom of the left hand-side N-terminal. The respective degrees of freedom of the right hand-side N-terminal do not feel a metadynamics potential. As a post-processing step, the initial 25 ns of each replica were excluded as equilibration, followed by the generation of the final structural ensemble by resampling the generated configurations based on the converged unbiasing weights. The converged unbiasing weights are obtained by following a previously reported procedure (37). To establish convergence, CVs as well as their time-dependent free energy profile for each CV are shown in **Figure S1B**. For molecular visualizations and calculating the local correlation of the final structural-ensemble generated cryo-EM map with the experimental cryo-EM map, we use Chimera and gmmconvert utility.

### Docking of aducanumab

The ColabFold notebook interface (61) (version 1.5.2) was utilized to generate the aducanumab model. The inputs included protein sequences and the alphafold2_multimer_v3 mode. The top-ranked model was selected as the final outcome. Subsequently, protein-protein docking between aducanumab and the most representative structure of both filament types was conducted using HADDOCK(62) (HADDOCK 2.4 server). For the filament, the N-terminal residues 1-8 and the core residues 18-21 were designated as active residues. The complementarity-determining regions (CDRs) of aducanumab (H1:31-35, H2:50-66, H3:99-113, and L1:21-34, L2:50-56, L3:89-97) were also chosen as active residues. The final complexes were selected taking in consideration the HADDOCK score and the geometric complementarity of the interfaces between the proteins. The poses were visually inspected to identify clashes, steric hindrances, and the quality of intermolecular interactions. Models that displayed favorable interactions, minimal steric clashes, and exhibited biologically relevant binding orientations were prioritized.

### Contact map, SASA and DSSP analysis

All the structural analysis are performed only on the two central chains of the fibrils, in order to avoid the finite size effects on the outermost chains. To calculate the contact map, perform SASA analysis, and analyse the secondary structure population (DSSP) the mdtraj (63) Python program was used. In the contact map analysis, a contact is considered to occur when the distance between the center of mass of two residues is less than 1 nm. The SASA analysis utilized the per-residue Shrake-Rupley function, with a selected probe radius of 0.34.

### Solvent diffusion analysis for water molecules and sodium ions

To calculate the diffusion coefficient of water molecules or sodium ions the mdanalysis toolkit was used (64). For each case, a cylinder centered at the middle chain along the filaments at residues E11 or H14/L21 or E22/D23 for type I filaments and G37 or I41/S26 or E22-D23 for type II filaments. Around these centers the radius of each cylinder was set to 1 nm and the height of the cylinder was chosen as 3.6 nm so that it covers the axis length of the filaments. Within these cylinders, the mean square deviation (MSD) of the oxygen atoms of water molecules or of the sodium ion was calculated for each of the three trajectories and average and standard deviations were obtained. From the averaged MSD, we used the Einstein equation to calculate the diffusion constant.

