## Supplementary Information for "Transient interactions between the fuzzy coat and the cross-β core of brain-derived Aβ42 filaments"

| Description | Group |
|---|---|
| RESIDUES=1-8 | tail0 |
| RESIDUES=85-92 | tail2 |
| RESIDUES=169-176 | tail4 |
| RESIDUES=253-260 | tail6 |
| RESIDUES=337-344 | tail8 |
| RESIDUES=421-428 | tail10 |
| RESIDUES=505-512 | tail12 |
| RESIDUES=589-596 | tail14 |
| RESIDUES=673-680 | tail16 |
| RESIDUES=757-764 | tail18 |
| RESIDUES=841-848 | tail20 |
| RESIDUES=925-932 | tail22 |
| RESIDUES=18-21 | epitope0 |
| RESIDUES=102-105 | epitope2 |
| RESIDUES=186-189 | epitope4 |
| RESIDUES=270-273 | epitope6 |
| RESIDUES=354-357 | epitope8 |
| RESIDUES=438-441 | epitope10 |
| RESIDUES=522-525 | epitope12 |
| RESIDUES=606-609 | epitope14 |
| RESIDUES=690-693 | epitope16 |
| RESIDUES=774-777 | epitope18 |
| RESIDUES=858-861 | epitope20 |
| RESIDUES=942-945 | epitope22 |

**Table S1**. List of residue groups used in the definition of the collective variables (CVs) used for biasing (**Table S2**).

| Description | Plumed file notation | Manuscript Notation |
|---|---|---|
| Coordination tail0 and tail2 | con0 | #C0 |
| Coordination tail0 and tail4 | con1 | #C1 |
| Coordination tail4 and tail22 | con2 | #C2 |
| Coordination tail20 and tail22 | con3 | #C3 |
| Coordination tail2 and tail6 | con4 | #C4 |
| Coordination tail6 and tail8 | con5 | #C5 |
| Coordination tail8 and tail14 | con6 | #C6 |
| Coordination tail10 and tail14 | con7 | #C7 |
| Coordination tail10 and tail12 | con8 | #C8 |
| Coordination tail12 and tail16 | con9 | #C9 |
| Coordination tail16 and tail18 | con10 | #C10 |
| Distance between the center of mass tail0 and epitope0 | dcm_0 | #d1 |
| Distance between the center of mass tail2 and epitope2 | dcm_2 | #d2 |
| Distance between the center of mass tail4 and epitope4 | dcm_4 | #d3 |
| Distance between the center of mass tail6 and epitope6 | dcm_6 | #d4 |
| Distance between the center of mass tail8 and epitope8 | dcm_8 | #d5 |
| Distance between the center of mass tail10 and epitope10 | dcm_10 | #d6 |
| Distance between the center of mass tail12 and epitope12 | dcm_12 | #d7 |
| Distance between the center of mass tail14 and epitope14 | dcm_14 | #d8 |
| Distance between the center of mass tail16 and epitope16 | dcm_16 | #d9 |
| Distance between the center of mass tail18 and epitope18 | dcm_18 | #d10 |
| Distance between the center of mass tail20 and epitope20 | dcm_20 | #d11 |
| Distance between the center of mass tail22 and epitope22 | dcm_22 | #d12 |
| alpha helical content tail0 | alpha0 | #ah1 |
| alpha helical content tail2 | alpha2 | #ah2 |
| alpha helical content tail4 | alpha4 | #ah3 |
| alpha helical content tail6 | alpha6 | #ah4 |
| alpha helical content tail8 | alpha8 | #ah5 |
| alpha helical content tail10 | alpha10 | #ah6 |
| alpha helical content tail12 | alpha12 | #ah7 |
| alpha helical content tail14 | alpha14 | #ah8 |
| alpha helical content tail16 | alpha16 | #ah9 |
| alpha helical content tail18 | alpha18 | #ah10 |
| alpha helical content tail20 | alpha20 | #ah11 |
| alpha helical content tail22 | alpha22 | #ah12 |
| parallel beta sheet content tail0 | parabeta0 | #bs1 |
| parallel beta sheet content tail2 | parabeta2 | #bs2 |
| parallel beta sheet content tail4 | parabeta4 | #bs3 |
| parallel beta sheet content tail6 | parabeta6 | #bs4 |
| parallel beta sheet content tail8 | parabeta8 | #bs5 |
| parallel beta sheet content tail10 | parabeta10 | #bs6 |
| parallel beta sheet content tail12 | parabeta12 | #bs7 |
| parallel beta sheet content tail14 | parabeta14 | #bs8 |
| parallel beta sheet content tail16 | parabeta16 | #bs9 |
| parallel beta sheet content tail18 | parabeta18 | #bs10 |
| parallel beta sheet content tail20 | parabeta20 | #bs11 |
| parallel beta sheet content tail22 | parabeta22 | #bs12 |

**Table S2**. List of collective variables (CVs) used for biasing.

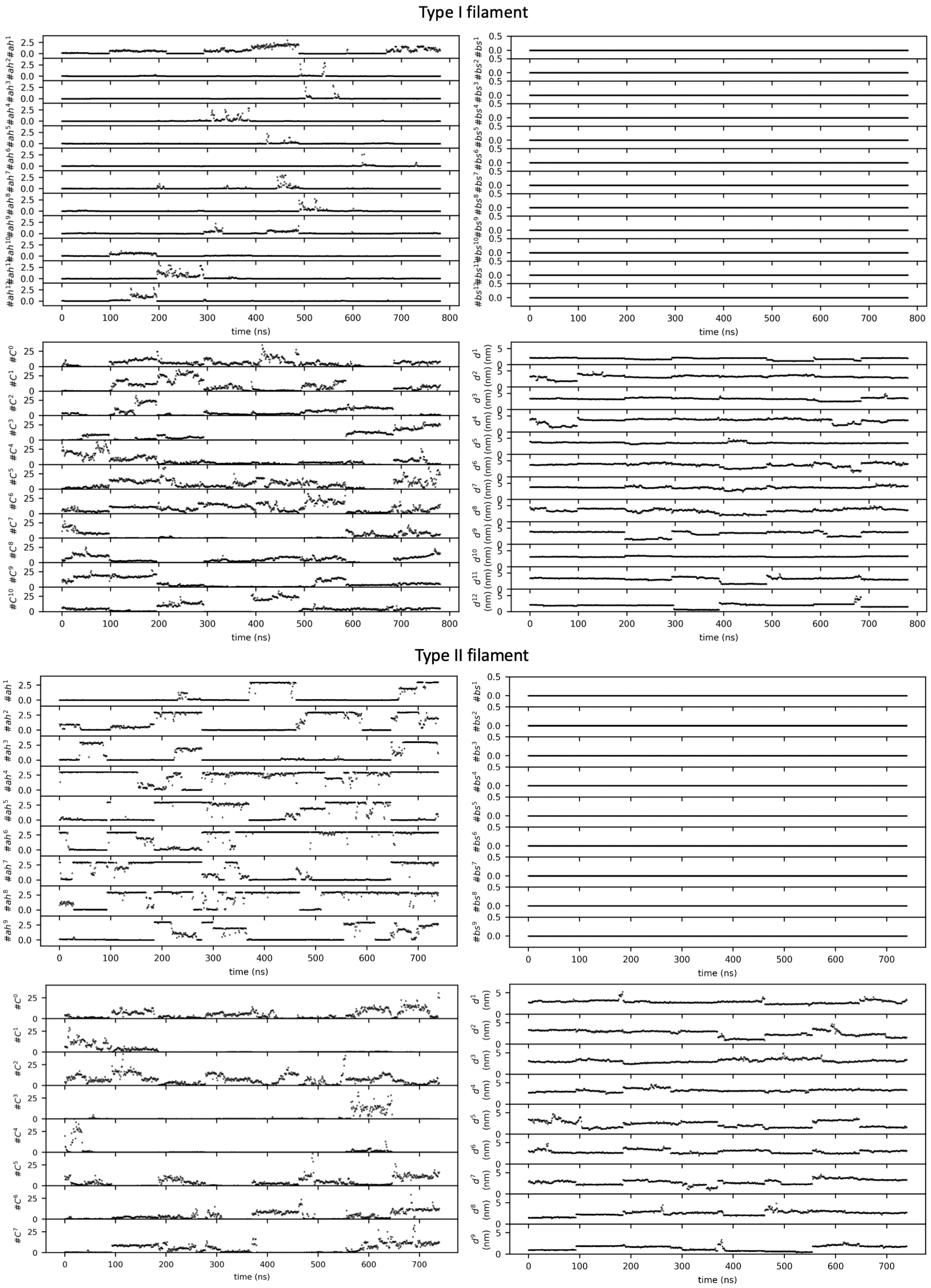


**Figure S1. Assessment of the convergence of the MEMMI simulations.** Time evolution profiles for all biased CVs for type I filament (up) and type II filament (down).

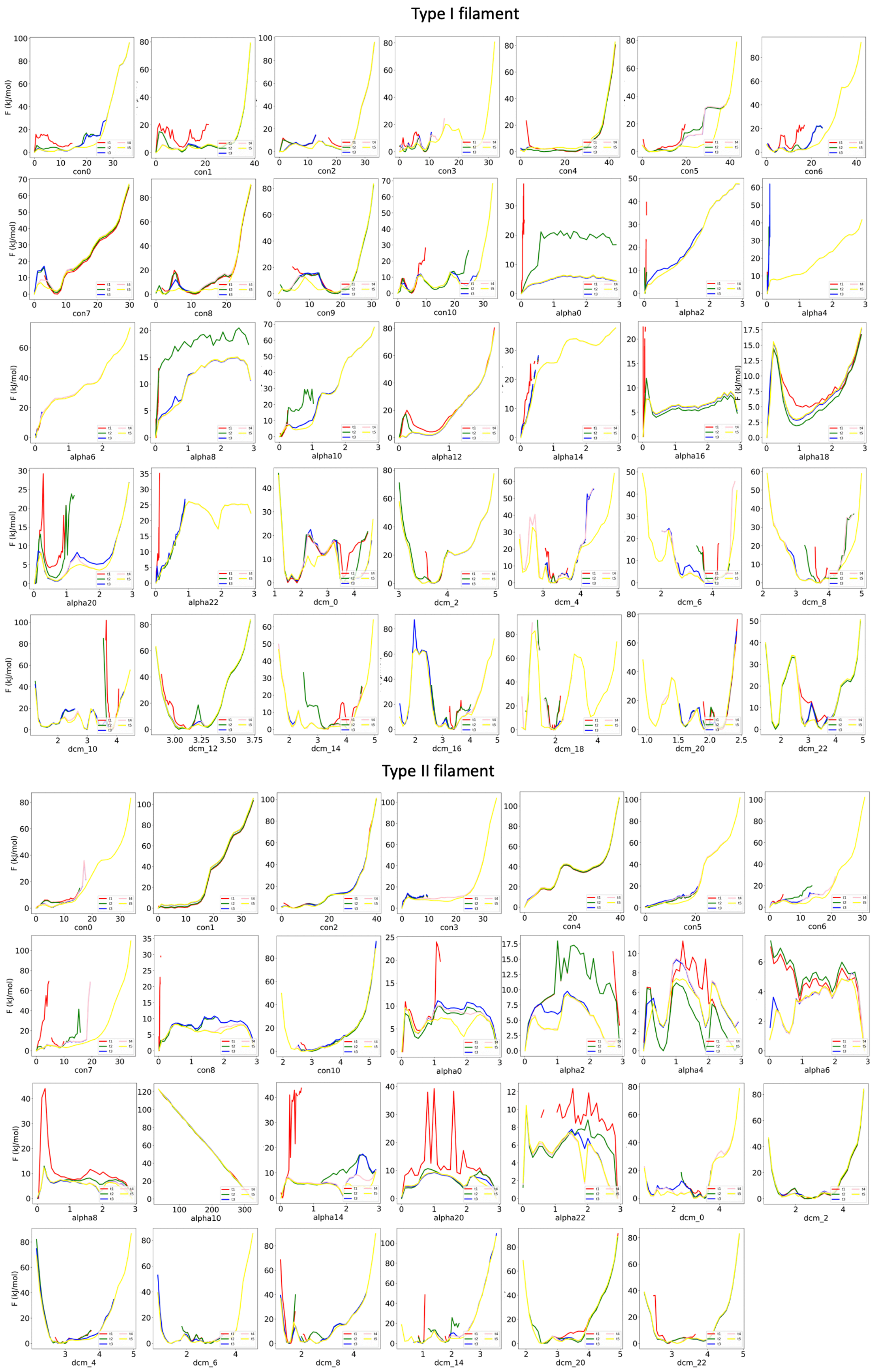


**Figure S2. Assessment of the convergence of the MEMMI simulations.** Free energy profiles for all biased CVs for subsequent 1 μs increments of simulated time.

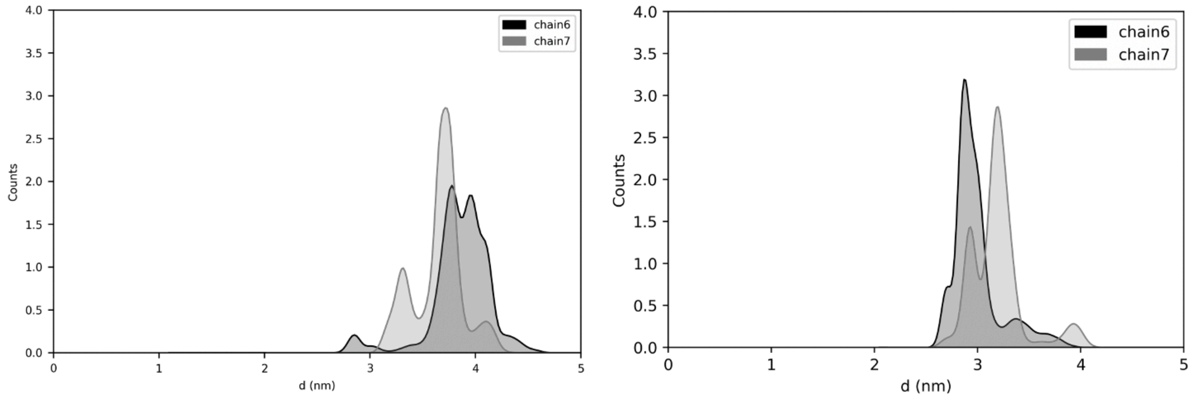


**Figure S3. Distance distribution between the fuzzy coat and the cross-β core of type I and type II filaments.** Distribution of the distance between the N-terminus (residues 1-9) and the cross-β core (residues 16-21) of the two central chains (chain 6 and chain 7) during the simulation. Type I filament (left panel) and type II filament (right panel).

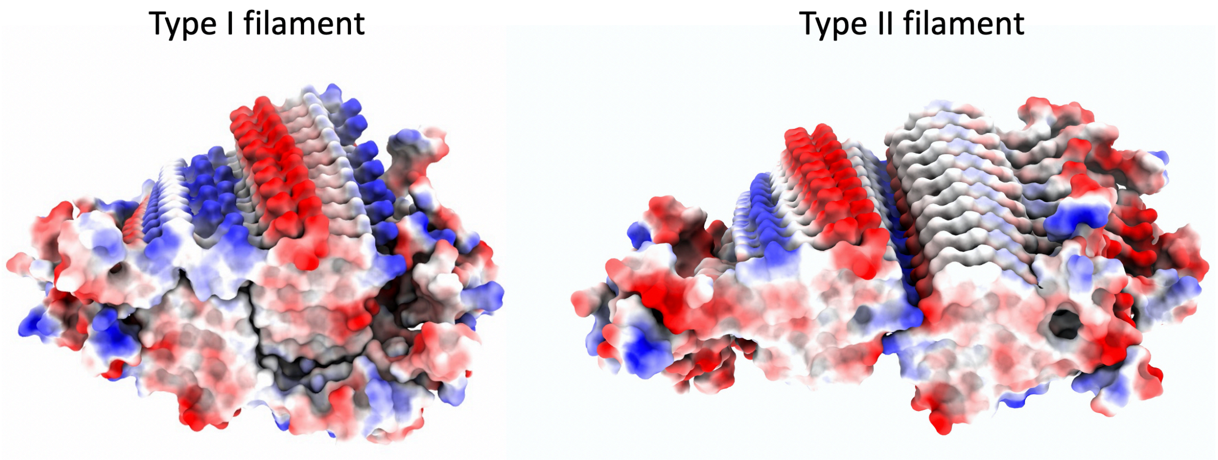


**Figure S4. Surface electrostatic potential of type I and type II filaments.** Most representative structure of the MEMMI structural ensemble of type I filament (left) and type II filament (right). Electrostatic potential maps are colored in blue to indicate positive potential and in red for negative potential.

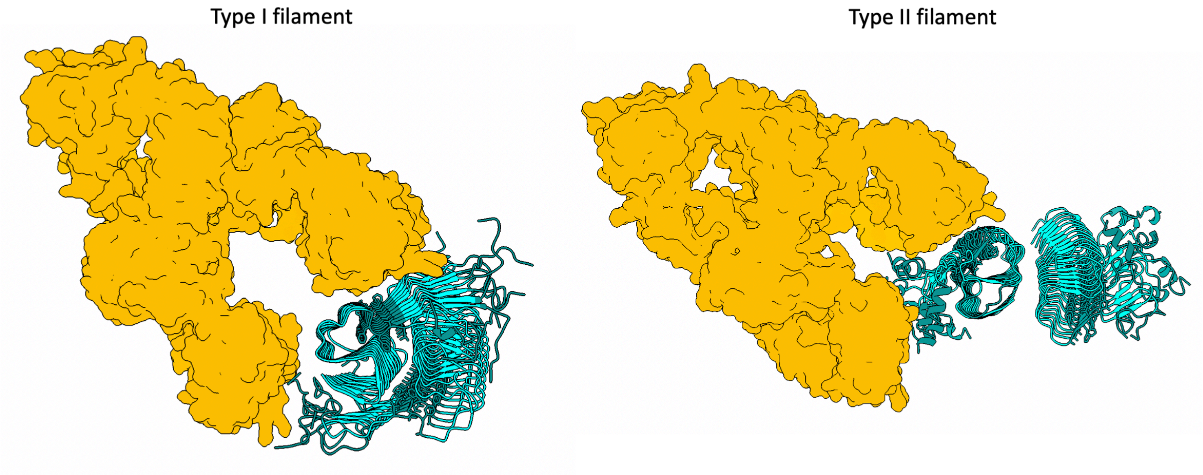


**Figure S5. Schematic representation of possible complexes between aducanumab and type I and type II filaments.** The cartoons show aducanumab in complex with type I (left panel) and type II (right panel) filaments, shown in orange with a surface representation. The cross-β core residues (16-21) and N-terminal residues (1-9) are highlighted in teal sticks.
